# Multidimensional Heritability Analysis of Neuroanatomical Shape

**DOI:** 10.1101/033407

**Authors:** Tian Ge, Martin Reuter, Anderson M. Winkler, Avram J. Holmes, Phil H. Lee, Lee S. Tirrell, Joshua L. Roffman, Randy L. Buckner, Jordan W. Smoller, Mert R. Sabuncu

**Author notes:** MRS and JWS contributed equally. Correspondance to (MRS) or (TG).

## Abstract

In the dawning era of large-scale biomedical data, multidimensional phenotype vectors will play an increasing role in examining the genetic underpinnings of brain features, behavior and disease. For example, shape measurements derived from brain MRI scans are multidimensional geometric descriptions of brain structure and provide an alternate class of phenotypes that remains largely unexplored in genetic studies. Here we extend the concept of heritability to multidimensional traits, and present the first comprehensive analysis of the heritability of neuroanatomical shape measurements across an ensemble of brain structures based on genome-wide SNP and MRI data from 1,320 unrelated, young and healthy individuals. We replicate our findings in an extended twin sample from the Human Connectome Project (HCP). Our results demonstrate that neuroanatomical shape can be significantly heritable, above and beyond volume, and can serve as a complementary phenotype to study the genetic determinants and clinical relevance of brain structure.

## Introduction

The exponential progress in genomic technologies has accelerated the examination of the genetic underpinnings of complex phenotypes, such as psychiatric and neurological disorders, many of which are highly heritable [1, 2]. For example, large-scale genome-wide association studies (GWAS) have provided insights about common genetic variants linked with a range of clinical conditions [3–6]. Most prior genetic studies have focused on univariate (scalar) phenotypes, such as diagnosis or a quantitative measurement. However, with the emergence of large-scale data collection efforts, such as the Human Connectome Project (HCP; http://www.humanconnectome.org) and the UK Biobank (http://www.ukbiobank.ac.uk), each subject can be linked to a high-dimensional phenotype vector, which might include imaging measurements or electronic health record. Such phenotypically rich studies open up the opportunity to analyze collections of multidimensional phenotypes, which can be more informative than scalar traits.

Brain imaging is playing an increasing role in the study of the relationship between genetic variants, neuroanatomy, behavior and disease susceptibility [7–10]. To date, most structural neuroimaging genetics studies have utilized the size, average cortical thickness, or surface area of a brain region to yield important discoveries about the genetic basis of brain morphology [see e.g., 11–14]. While these measurements capture a few basic dimensions of anatomical variability, they provide a limited description of the underlying geometry.

Neuroanatomical shape measurements — multidimensional geometric descriptions of brain structure — have attracted increasing attention in medical image analysis. Shape measurements characterize isometry-invariant (in particular, independent of location and orientation) geometric attributes of an object, which provide a rich description of an anatomical structure and can encompass volumetric variation. Such measurements may thus offer increased sensitivity and specificity in examining the clinical relevance and genetic underpinnings of brain structure. Recent studies have shown that the shape of subcortical brain regions and cortical folding patterns provide information that is not available in volumetric measurements and is predictive of disease status, onset and progression in schizophrenia [15–17], autism [18, 19], bipolar disorder [20, 21], Alzheimer’s disease [22–25], and other mental disorders [26, 27]. There is also increasing evidence that genetic variants may have influences on brain morphology that can be captured by shape measurements [28–32].

This paper makes two major contributions to the investigation of the genetic basis of neuroanatomical shape. First, we extend the theoretical concept of heritability to multidimensional traits, such as the shape descriptor of an object, and propose a novel method to estimate the heritability of multidimensional traits based on genome-wide single nucleotide polymorphism (SNP) data from unrelated individuals (known as SNP heritability). Our estimation method builds on genome-wide complex trait analysis (GCTA) [33, 34] and phenotype correlation-genetic correlation (PCGC) regression [35], and generalizes these techniques to the multivariate setting. Second, using structural magnetic resonance imaging (MRI) and SNP data from 1,320 unrelated individuals collected as part of the Harvard/Massachusetts General Hospital (MGH) Brain Genomics Superstruct Project (GSP) [36, 37], we present the first comprehensive heritability analysis of the shape of an ensemble of brain structures, quantified by the truncated Laplace-Beltrami Spectrum (LBS) (also known as the “Shape-DNA”) [38–40], in this young (18-35 years) and healthy cohort, and devise a strategy to visualize primary modes of shape variation. We also replicate our findings in an extended twin sample (72 monozygotic twin pairs, 69 dizygotic twin pairs, 253 full siblings of twins and 55 singletons) from the Human Connectome Project (HCP) [41].

The truncated LBS is a multidimensional shape descriptor, which can be obtained by solving an eigenvalue problem on the 2D boundary surface representation of an object. It is invariant to the representation of the object including parameterization, location and orientation, and thus does not require spatial alignment with a population template, making it computationally efficient and robust to registration errors. LBS also depends continuously on topology-preserving deformations, and is thus suitable to quantify differences between shapes. Recent empirical evidence suggests that the LBS-based shape descriptor provides a discriminative characterization of brain anatomy and offers state-of-the-art performance for a range of shape retrieval and segmentation applications [42, 43]. A collection of the descriptors of brain structures, known as the BrainPrint, can provide an accurate and holistic representation of brain morphology, and has been successfully applied to subject identification, sex and age prediction, brain asymmetry analysis, twin studies, and computer-aided diagnosis of dementia [40, 44]. Our LBS-based heritability analyses demonstrate that neuroanatomical shape can be significantly heritable, above and beyond volume, and yield a complementary phenotype that offers a unique perspective in studying the genetic determinants of brain structure.

## Results

### Heritability of the volume of neuroanatomical structures

To benchmark our shape results, we first computed SNP heritability estimates for the volumetric measurements of an array of brain regions using the GSP sample. Table 1 lists these heritability estimates after adjusting for intracranial volume (ICV or head size) as a covariate. Point estimates of the heritability of volumetric measurements suggested that several neuroanatomical structures have moderately heritable volumes. In particular, the caudate, corpus callosum, lateral ventricle, 3rd and 4th ventricles, pallidum, putamen and thalamus all had volume heritability estimates greater than 25%. Table 1 further includes p-values for the statistical significance of the heritability estimates. The parametric (Wald) and non-parametric (permutation-based) p-values were virtually identical, confirming the accuracy of the standard error estimates we computed (see Methods). We observe that none of the volume heritability estimates were statistically significant after correcting for multiple comparisons (false discovery rate or FDR at q = 0.05), possibly due to sample size limitations. Only the volumes of the caudate, corpus callosum and 3rd ventricle achieved a heritability that was nominally significant in our sample (uncorrected p < 0.05). Table 1 also includes test-retest reliability estimates of volume after regressing out ICV, computed as Lin’s concordance correlation coefficient [45] using measurements from 42 subjects with repeated scans on separate days. Almost all the structures had a volume estimate reliability greater than 0.75 except for the pallidum. There was no significant correlation between the reliability and heritability estimates of volume (p = 0.828).

**Table 1:** SNP heritability estimates 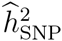 of the volume of brain structures using the GSP sample. The standard errors (SE) were computed using an approximation, which, given the empirical genetic similarity matrix, only depends on the sample size. *p*-values were obtained by the Wald test and the permutation inference (based on 10,000 permutations), respectively. The strong agreement between the parametric and non-parametric *p*-values indicates that the estimated SE values are accurate. Estimates with uncorrected significant *p*-values (< 0.05) are shown in **bold**. Test-retest reliability of the volumetric measurements was computed as Lin’s concordance correlation coefficient using measurements from 42 subjects with repeated scans on separate days.

| Structure | $\hat{h}_{\text{SNP}}^2$ | SE | Wald $p$-value | Perm $p$-value | Reliability |
| --- | --- | --- | --- | --- | --- |
| Accumbens Area | 0.001 | 0.281 | 0.500 | 1.000 | 0.797 |
| Amygdala | 0.141 | 0.281 | 0.308 | 0.305 | 0.864 |
| Caudate | **0.657** | 0.281 | 0.010 | 0.009 | 0.947 |
| Cerebellum | 0.084 | 0.281 | 0.383 | 0.382 | 0.989 |
| Corpus Callosum | **0.538** | 0.281 | 0.028 | 0.029 | 0.882 |
| Hippocampus | 0.005 | 0.281 | 0.493 | 0.492 | 0.939 |
| Lateral Ventricle | 0.331 | 0.281 | 0.119 | 0.120 | 0.995 |
| 3rd Ventricle | **0.500** | 0.281 | 0.038 | 0.040 | 0.832 |
| 4th Ventricle | 0.381 | 0.281 | 0.087 | 0.089 | 0.986 |
| Pallidum | 0.300 | 0.281 | 0.142 | 0.142 | 0.642 |
| Putamen | 0.328 | 0.281 | 0.121 | 0.122 | 0.934 |
| Thalamus | 0.252 | 0.281 | 0.184 | 0.186 | 0.867 |

### Heritability of the shape of neuroanatomical structures

Neuroanatomical shape measurements provide a geometric characterization and a rich description of brain structure. We therefore hypothesize that analyzing the shape variation of neuroanatomical structures can identify genetic influences beyond captured by volumetric measurements. Fig. 1 and Table 2 show the SNP heritability estimates of the shape of an ensemble of brain structures using the GSP sample. These estimates were computed based on LBS descriptors normalized for size and explicitly including the volume of the corresponding structure as a covariate in the analysis to account for potential volume effects. A number of structures showed moderate to high SNP heritability. Specifically, the shape of the caudate, cerebellum, corpus callosum, hippocampus, 3rd ventricle and putamen exhibited heritability estimates greater than 25%. All these estimates were statistically significant after correcting for an FDR at *q* = 0.05. We observe that this is in contrast with the case of volume, where despite a similar heritability range, no estimate reached FDR-corrected significance. The main reason for this discrepancy is the theoretically guaranteed reduced standard errors in SNP heritability estimates of multidimensional traits (see Methods for a theoretical treatment). The shape of the accumbens area was also marginally significantly heritable with an uncorrected *p*-value less than 0.05. As in the case of volume, the parametric (Wald) *p*-values were virtually identical to the permutation *p*-values, suggesting that our standard error estimates are accurate (see Methods).

**Figure 1:**
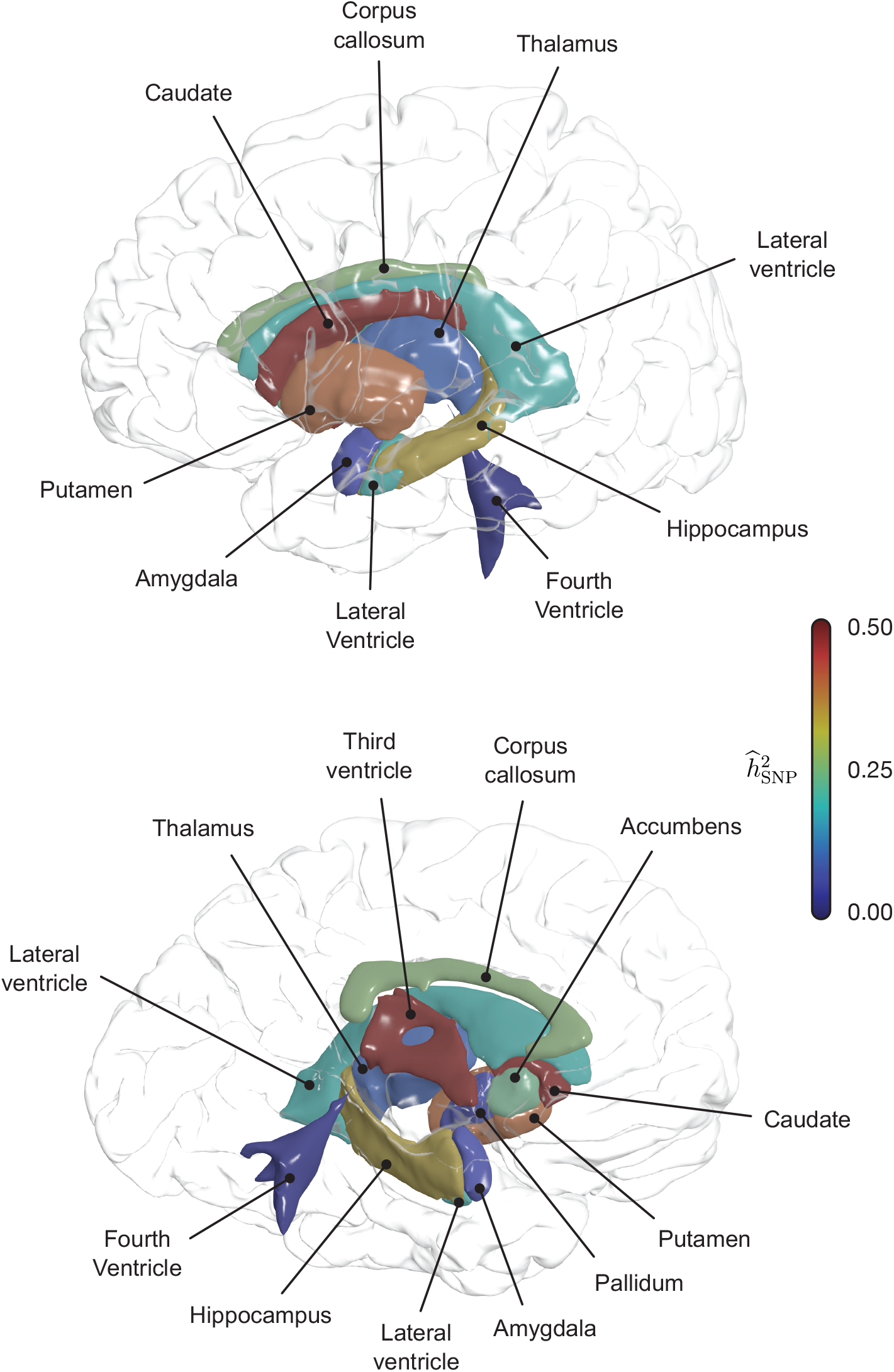
SNP heritability estimates of the shape of brain structures in the GSP sample. Top: lateral view. Bottom: medial cross-section.

**Table 2:** SNP heritability estimates 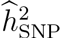 of the shape of brain structures using the GSP sample. Standard errors (SE) are less than those corresponding to volume heritability. *p*-values were obtained by the Wald test and the permutation inference (based on 10,000 permutations), respectively. The strong agreement between the parametric and non-parametric *p*-values indicates that the SE estimates are accurate. Estimates with uncorrected significant *p*-values (< 0.05) are shown in bold. False discovery rate (FDR) corrected significant *p*-values (< 0.05) are shown in red. Test-retest reliability of the shape measurements were computed as the average Lin’s concordance correlation coefficient of individual components of the LBS-based shape descriptor from 42 subjects with repeated scans on separate days.

| Structure | $\hat{h}_{\text{SNP}}^2$ | SE | Wald $p$-value | Perm $p$-value | Reliability |
| --- | --- | --- | --- | --- | --- |
| Accumbens Area | **0.237** | 0.135 | 0.039 | 0.039 | 0.418 |
| Amygdala | 0.061 | 0.139 | 0.330 | 0.327 | 0.670 |
| Caudate | **0.499** | 0.188 | 0.004 | 0.005 | 0.759 |
| Cerebellum | **0.452** | 0.192 | 0.009 | 0.009 | 0.844 |
| Corpus Callosum | **0.264** | 0.133 | 0.023 | 0.022 | 0.622 |
| Hippocampus | **0.347** | 0.169 | 0.020 | 0.019 | 0.866 |
| Lateral Ventricle | 0.190 | 0.153 | 0.107 | 0.105 | 0.890 |
| 3rd Ventricle | **0.500** | 0.157 | 0.001 | 0.001 | 0.761 |
| 4th Ventricle | 0.005 | 0.208 | 0.490 | 0.491 | 0.633 |
| Pallidum | 0.061 | 0.117 | 0.299 | 0.299 | 0.402 |
| Putamen | **0.413** | 0.148 | 0.003 | 0.003 | 0.781 |
| Thalamus Proper | 0.086 | 0.143 | 0.274 | 0.276 | 0.552 |

Table 2 also lists test-retest reliability estimates for the shape of different structures. Analogous to the case of volume, we quantified reliability as the average Lin’s concordance correlation coefficient of individual components of the multidimensional shape descriptor from 42 subjects with repeat scans on separate days. These results suggest that the LBS-based shape descriptors were overall less reliable than volumetric measurements, with half of the structures exhibiting a shape reliability less than 0.75. This is likely due to the increased sensitivity of shape to segmentation differences relative to the volume. Furthermore, there was a marginally significant correlation between reliability and heritability of shape (Pearson’s *r* = 0.562 and *p* = 0.057). We conclude that close to 30% of the variation in shape heritability across structures can be attributed to the reliability of the shape descriptor. This suggests that for structures that exhibited low shape heritability (e.g., amygdala), a more accurate image segmentation and shape analysis pipeline might yield an increased estimate of heritability. We further conducted a sensitivity analysis of shape heritability estimates, with respect to the two free parameters of the LBS-based shape descriptor: number of eigenvalues incorporated and amount of smoothing applied to the surface mesh representing the geometry of the object. Supplementary Fig. 1 shows that the heritability estimates were largely robust to variations in these parameters.

We sought to replicate these findings in the HCP sample, in which subjects are healthy and have a similar age range as the GSP sample. We selected 590 non-Hispanic/Latino Europeans aged between 22 and 35, comprising 72 monozygotic (MZ) twin pairs, 69 dizygotic (DZ) twin pairs, 253 full siblings of twins and 55 singletons (single birth individuals without siblings). We estimated the shape heritability for brain structures that had significantly heritable shapes in the GSP sample using an ACE model (A: additive genetics; C: common environment; E: unique or subject-specific environment), where the additive genetic similarity was derived from pedigree information and the common environment term reflected household sharing between subjects. We also obtained the standard error of the shape heritability estimates using a block bootstrapping procedure (see Methods). All the shapes we analyzed were significantly heritable in the HCP sample: accumbens area 0.309 ± 0.081; caudate 0.583 ± 0.062; cerebellum 0.653 ± 0.060; corpus callosum 0.558 ± 0.068; hippocampus 0.363 ± 0.095; 3rd ventricle 0.536 ± 0.067; putamen 0.483 ± 0.106. We also observe that the HCP shape heritability estimates were consistently larger than the GSP estimates, which is theoretically expected because the SNP heritability estimated from unrelated GSP subjects only captured the genetic variation tagged by common SNPs in the data set, and is thus a lower bound for the classical narrow-sense heritability estimated from familial data such as the HCP sample.

### Visualizing the principal mode of shape variation

The LBS-based shape descriptor is suitable to efficiently and accurately extract intrinsic properties of the shape of brain structures from a large number of individuals, but is not designed to visually inspect shape differences. Here, we propose a strategy to visualize the principal mode of shape variation. Specifically, it can be shown that the first principal component (PC) of the multidimensional LBS-based shape descriptor captures the greatest shape variation and has the largest impact on the overall heritability estimate of the shape (see Methods). We thus visualized shape variation along the first PC of the shape descriptor for brain structures with significantly heritable shapes in the GSP sample: right caudate, cerebellum, corpus callosum, right hippocampus, 3rd ventricle, and left putamen. The illustrations of contralateral structures (i.e., left caudate, left hippocampus and right putamen), which showed similar shape variation, are provided in Supplementary Fig. 2. In each panel of Fig. 2, the structure is represented with a sample-specific population average, on which average shapes at the two extremes (±2 standard deviation or SD) of the principal axis with identical volume (−2 SD, blue; +2 SD, red) are depicted. Blue regions indicate where shapes around the −2 SD are larger than shapes around the +2 SD, and vice versa for the red regions.

**Figure 2:**
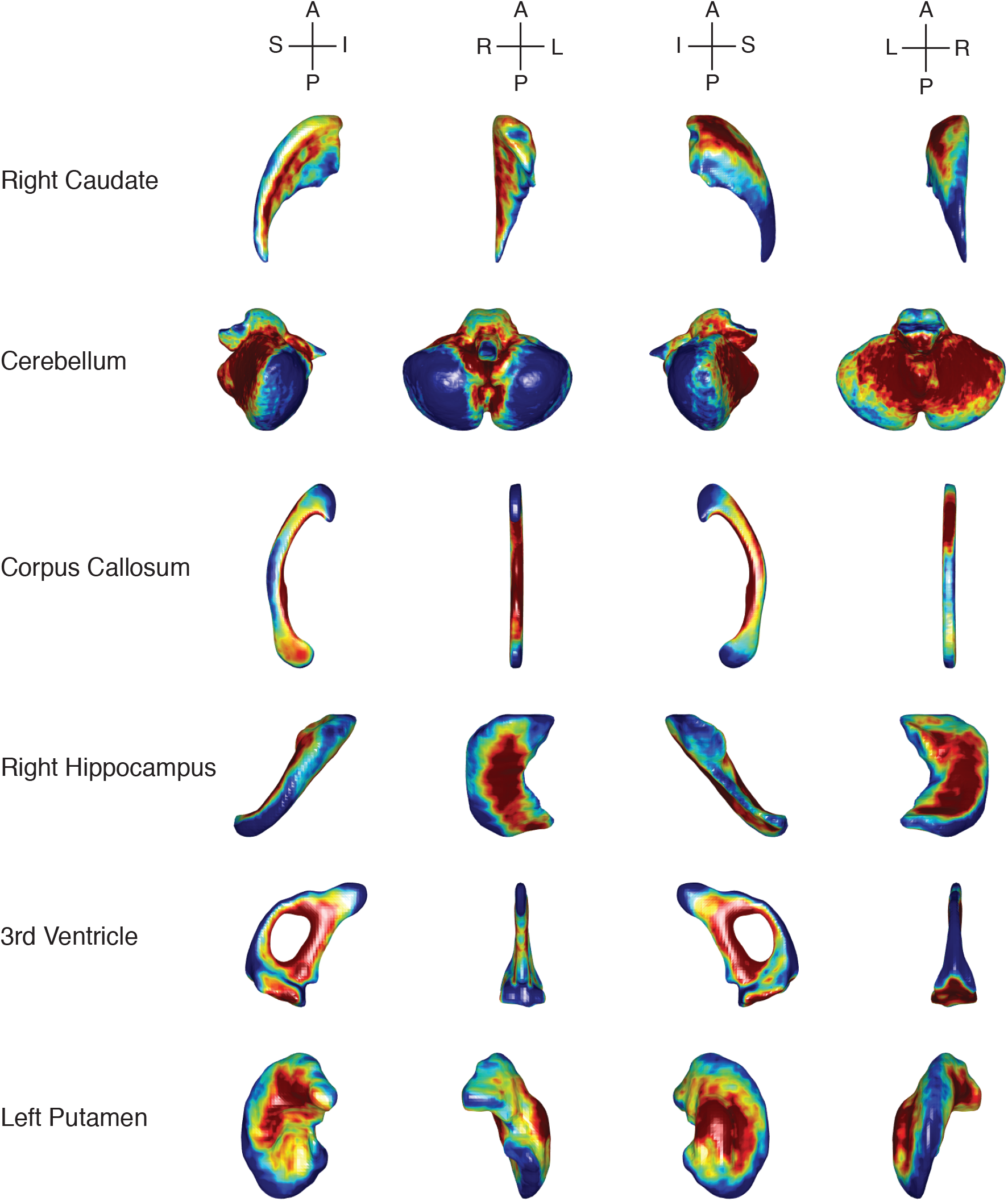
The principal mode of shape variation for brain structures with significantly heritable shape in the GSP sample. Each structure is represented with a sample-specific population average, on which average shapes at the two extremes (±2 standard deviation or SD) along the first principal component (PC) of the shape descriptor (−2 SD, blue; +2 SD, red) are depicted. Anatomical orientation is indicated with embedded coordinate axes. I: Inferior, S: Superior, A: Anterior, P: Posterior, L: Left, R: Right.

The first PC of the right caudate captured 77% of the shape variation and had a SNP heritability estimate of 0.88. Moving along the principal mode of shape variation, the right caudate had a larger (smaller) head with a corresponding shorter (longer) tail. For the cerebellum, the first PC explained 69% of the shape variation and had a SNP heritability of 0.61. A clear expansion (contraction) of the anterior lobe and a corresponding contraction (expansion) of the posterior lobe can be observed along the principal axis. The first PC of the corpus callosum captured 41% of the shape variation and had a SNP heritability of 0.41. The principal mode captured an expansion (contraction) of the middle corpus callosum along the dorsoventral axis and a corresponding shrinking (enlargement) of the anterior and posterior part of the structure. For the right hippocampus, the first PC explained 69% of the shape variation, had a SNP heritability of 0.47, and exhibited dorsoventral widening (narrowing) of the body and corresponding lateral and anterior-posterior contraction (expansion). The first PC of the 3rd ventricle captured 69% of the shape variation and had a SNP heritability estimate of 0.80. The principal mode captured an enlargement (shrinking) of the posterior protrusions and an expansion (contraction) of the lateral walls, coupled with a corresponding contraction (expansion) of the roof of the cleft. Finally, first PC of the left putamen explained 61% of the shape variation, had a SNP heritability of 0.70, and captured lateral widening (narrowing) and a corresponding contraction (expansion) along the dorsoventral and anterior-posterior axes.

## Discussion

This work makes two contributions to neuroscience and genetic research. First, we extend the concept of heritability to multidimensional traits and present an analytic strategy that generalizes SNP heritability analysis. The heritability estimator we propose for multidimensional traits has reduced uncertainty in its point estimate relative to univariate estimates, and thus offers more statistical power. Our empirical analyses confirmed this theoretical expectation. Moreover, and in the same line, we provide methods that can easily adjust for covariates in multivariate models, and also both parametric and nonparametric inferential tools that can assess the significance of a heritability estimate. Our approach opens the door to the genetic characterization of shape measurements and other multidimensional traits.

Second, we use the proposed approach to quantify the SNP heritability of the shape of an ensemble of anatomical structures spanning the human brain in a group of young and healthy subjects. The shape of caudate, cerebellum, corpus callosum, hippocampus, 3rd ventricle and putamen exhibited moderate to high heritability (i.e., greater than 25%), after controlling for volume. All of these estimates achieved FDR-corrected significance at *q* = 0.05. This is in contrast to the volume heritability estimates of the same set of brain structures on the same sample, none of which reached FDR-corrected significance. Although our heritability analysis of volume, which was used to benchmark the shape analysis, may be less informative compared to more powerful twin studies and large-scale meta-analyses in the literature, the increased statistical power and the additional information shape analysis can provide relative to volumetric analysis demonstrate the usefulness of our methods and underscore the potential of leveraging multidimensional traits when analyzing data sets with moderate sample sizes.

Using the extended twin data from the HCP, we also replicated significant shape heritability estimates observed in the GSP sample. Our HCP estimates were consistently larger than the GSP estimates, which is theoretically expected because SNP heritability estimated from the unrelated sample in GSP does not capture genetic contributions (e.g., from rare variants) that are not tagged by genotyped SNPs, and thus explains a smaller proportion of the phenotypic variation. However, additional factors may contribute to the difference between SNP and familial heritability estimates, which include improper modeling of shared environment, assortative mating, genetic interaction (epistasis), suboptimal statistical methods for heritability estimation, and differences in sample characteristics such as age range, ethnic background and environmental exposures [35, 46–51]. Dissecting the discrepancy in heritability estimates from familial and unrelated data is an area under active investigation. More systematic future work is required to fully disentangle this problem.

A handful of prior neuroimaging studies have explored the shape of certain brain structures as potential phenotypes in examining genetic associations. For example, Qiu et al. [28] and Shi et al. [29] reported influences of the apolipoprotein E (APOE) *ε*4 allele on hippocampal morphology in depressive and Alzheimer’s disease patients. Variants involved in the regulation of the *FKBP5* gene were recently associated with hippocampal shape [30]. A meta-study [32] identified a GWAS significant SNP that exerts its effect on the shape of putamen bilaterally. Prior studies have also estimated heritability of shape based on familial relatedness. In a recent study, the heritability of the shape of subcortical and limbic structures was estimated using data from multigenerational families with schizophrenia [31]. In other related work, Mamah et al. [52] and Harms et al. [53] revealed shape abnormalities in basal ganglia structures (caudate, putamen and globus pallidus) and the thalamus in siblings of schizophrenia patients. An application of the LBS-based shape descriptor to twin data found increased shape similarity of brain structures in MZ twin pairs relative to DZ twins, indicating genetic influences on brain morphology [40], although heritability was not estimated.

However, to date, outside of these notable exceptions, most structural imaging genetics studies have utilized scalar measurements (e.g., volume, thickness, area) as phenotypes. In the present study, we accounted for potential volume effects in our shape analyses by normalizing the LBS-based shape descriptor for size and additionally including the volumetric measurement of the corresponding structure as a covariate when estimating heritability. Our results show that shape measurements provide a rich and novel set of phenotypes for exploring the genetic basis of brain structure, and may identify novel genetic influences on the brain that are not detectable with conventional analyses based on the volume of structures.

There are several biological mechanisms that might lead to shape differences with minimal effect on the overall size of the structure. These include localized volumetric effects that are confined to subfields, subnuclei or other sub-regions that make up the structure. Shape analysis may provide significant information about neurodevelopmental abnormalities, such as those associated with neuronal migration, synaptogenesis, synaptic pruning and myelination. Shape measurements might for example shed light on morphogenetic mechanisms that involve mechanical tensions along axons, dendrites and glial processes [54]. Thus, shape measurements are particularly promising phenotypes for studying neurodevelopmental disorders. Neurodegenerative processes and other pathologies, many of which are known to be genetically influenced, can also impact neuroanatomical shape by exerting focal and/or selective insults. For example, in Alzheimer’s disease, morphological alterations in the hippocampus may only target certain subfields [55].

The shape analysis literature offers an expanding list of methods to quantify and characterize shape [43]. A major advantage of the LBS-based shape descriptor [38] employed in this study is that it is robust to intensity variation across scans and does not require the nonlinear spatial registration of the object with a population template, which can be computationally demanding and prone to error. In this paper, we also present a novel strategy to visualize the principal mode of shape variation across the population. For brain structures with significantly heritable shapes, we demonstrated that the principal mode explains a large portion of the overall shape variation and is often highly heritable. This approach can thus shed light on the global genetic influences on brain structures, and is complementary to studies that rely on nonlinear group-wise registration to characterize localized genetic influences on shape variation.

In the present study, in light of the similar shape variation of bilateral brain structures as observed in Fig. 2 and Supplementary Fig. 2, and to increase signal-to-noise ratio and statistical power, we combined the left and right structures in our shape heritability analysis. SNP heritability estimates of the shape of bilateral brain structures using the GSP sample are provided in Supplementary Table 1. The results indicate that the genetic influences on several anatomical structures (e.g., caudate) may be lateralized, although with the current sample size we are not able to claim that the lateral difference is significant. It will be interesting to investigate the laterality of brain structures from the genetic perspective when we have a better understanding of the genetic basis of brain morphology and when a data set with larger sample size becomes available.

The heritability analysis of multidimensional traits developed here can be applied to phenotypes other than shape that are intrinsically multivariate. Another application might involve heritability or genetic association analyses combining related traits to obtain more stable effect estimates. For example, it can be used as an alternative to principal component analysis (PCA) and factor analysis when investigating the genetic basis of various psychometric or behavioral traits. Also, voxel- or vertex-level neuroimaging measurements are often noisy, and analyzing these measurements in homogeneous brain regions in a multivariate fashion may increase the reliability and reproducibility of the results. Finally, the empirical genetic similarity matrix can be computed with other SNP grouping strategies (e.g., based on genes, pathways, functional annotations and previous GWAS findings) to model the genetic influences from a specific genomic region or partition the heritability of multidimensional traits, as in Yang et al. [56].

## Methods

### Variance component models

We start with a brief review of variance component models (also known as random effects models) for the heritability analysis of univariate (scalar) traits, which provide a general statistical framework that can handle both familial designs and unrelated individuals randomly sampled from the population. Assuming, for the moment, no covariate needs to be adjusted, and a trait can be partitioned into the sum of additive genetic effect *g*, common (or shared) environment *c*, and unique (subject-specific) environment *e*, the variance component model takes the following form:

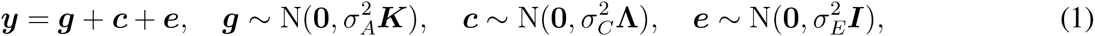

where *y* = [*y*_1_, …, *y*_*N*_]^T^ is a vector comprising quantitative traits from *N* individuals, 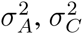 and 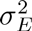 are the additive genetic variance, common environmental variance and unique environmental variance, respectively, *K* is a genetic similarity matrix, Λ quantifies shared environment between pairs of individuals, and *I* is an identity matrix.

In familial studies, *K* is twice the kinship matrix, *K* = 2Φ, and indicates expected additive genetic covariance among individuals. The *ij*-th entry of the kinship matrix, *ϕ*_*ij*_, known as the kinship coefficient, defines genetic relatedness for subjects *i* and *j*, and in general can be derived from pedigree information [57, 58]. Λ is a matrix that usually reflects household sharing between pairs of individuals. For example, in the present study, *ϕ*_*ij*_ = 1/2 for MZ twins and *ϕ*_*ij*_ = 1/4 for DZ twins and full siblings, and we assume that twin pairs and their non-twin siblings share the same environment and the corresponding elements in Λ are 1.

When modeling unrelated individuals randomly sampled from the population, *K* is the empirical genetic similarity matrix for each pair of individuals estimated from genome-wide SNP data, and the corresponding variance component parameter 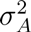 is the total additive genetic variance tagged by common SNPs spanning the genome. We note that in unrelated subject studies 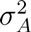 does not capture contributions (e.g., from rare variants) that are not assayed by the genotyping microarray, and thus needs to be interpreted differently from 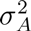 in familial studies, although we use the same notation here for simplicity. In addition, the common environmental matrix Λ is often assumed to vanish when analyzing unrelated individuals, in which case Eq. (1) becomes the classical model used in genome-wide complex trait analysis (GCTA) [33, 34].

The heritability of a univariate (scalar) trait is defined as

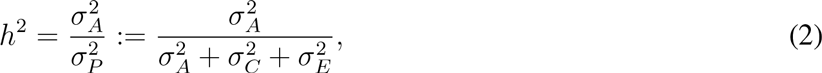

where 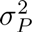 is the phenotypic variance. In familial studies, *h*^2^ measures the narrow-sense heritability of a trait, while in unrelated subject studies, *h*^2^ measures additive heritability attributable to common genetic variants (known as SNP heritability and often denoted as 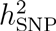), and provides a lower bound for the narrow-sense heritability estimated by familial studies.

### Heritability of multidimensional traits

We now consider an *M*-dimensional trait ***Y*** = [*y*_1_, …, *y*_*M*_] = [*y*_*im*_]_*N×M*_. We model ***Y*** by a multivariate variance component model:

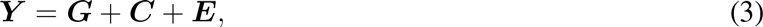

where ***G, C*** and ***E*** are *N* × *M* matrices, and represent additive genetic effects, common environmental factors and unique environmental factors, respectively. We have the following distributional assumptions:

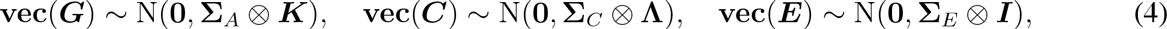

where vec(·) is the matrix vectorization operator which converts a matrix into a vector by stacking its columns, ⊗ is the Kronecker product of matrices, Σ*_A_* = [σ_*A*_*rs*__]_*M×M*_ is the genetic covariance matrix, whose *rs*-th element *σ*_*A*_*rs*__ is the genetic covariance between *y*_*r*_ and *y*_*s*_, Σ_*C*_ = [σ_*C*_*rs*__]_*M×M*_ is the common environmental covariance matrix, and ΣE = [σ_Ers_]MxM is the unique environmental covariance matrix. The distributional assumptions in Eq. (4) indicate that both the genetic effects and the environmental factors can be correlated across trait dimensions. When the trait is a scalar, the multivariate model (3) degenerates to the univariate model specified in Eq. (1). Analogous to the discussion above, *K* is derived from pedigree information in familial studies and empirically estimated from genome-wide SNP data in unrelated subject studies. As a result, Σ*_A_* denotes the genetic covariance due to common SNPs when analyzing unrelated subjects.

We define the heritability of a multidimensional trait as

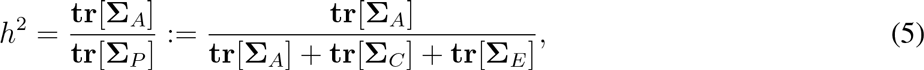

where Σ_*P*_ = [σ*_P_rs__*]*_M×M_* is the phenotypic covariance matrix and **tr**[·] is the trace operator of a matrix. This definition measures the proportion of the total phenotypic variance tr[Σ_P_] that can be explained by the total additive genetic variance tr[Σ_*A*_], and yields a heritability measure that is bounded between 0 and 1. When the trait is univariate, Σ*_A_*, Σ_*C*_, and Σ_*E*_ become scalars, and Eq. (5) reduces to the classical definition of heritability in Eq. (2). We use 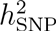 in place of *h*^2^ in unrelated subject studies to emphasize that it only captures genetic influences due to common genetic variants and is a lower bound for the narrow-sense heritability estimated using familial designs.

### Properties of multidimensional heritability

Our definition of heritability is invariant to rotations of the data. For a linear transformation ***T*** applied to the trait dimensions in model (3), i.e.,

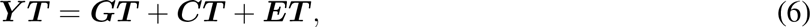

using the properties of vectorization and the Kronecker product, the covariance structure of the transformed trait ***YT*** can be computed as follows:

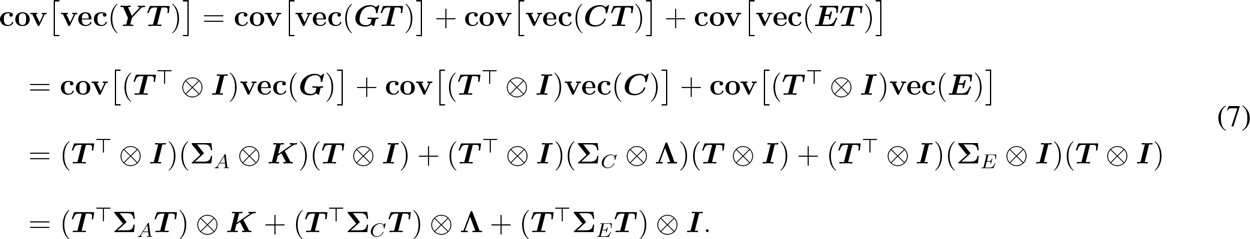

Therefore, the transformed heritability is

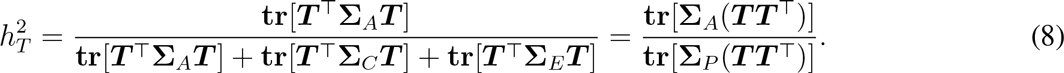

When *T* is an orthogonal matrix satisfying *TT*^T^ = *T*^T^*T* = *I*, we have 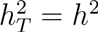.

The definition of heritability in Eq. (5) can also been written as

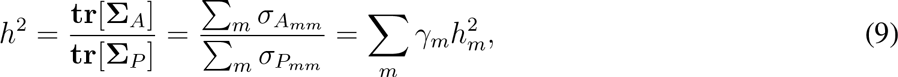

where *γ*_*m*_ = σ_*P*_*mm*__/Σ_*m*_ σ_*P*_*mm*__ with Σ*_m_*γ*_m_* = 1, and 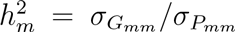 is the heritability of the *m*-th component of the trait. Therefore, our definition of the heritability of a multidimensional trait is essentially a weighted average of the heritability of its individual components.

### A moment-matching estimator for unrelated subject studies

The model (3) can in principle be fitted using likelihood-based methods. However, this can be computationally expensive when the dimension of the trait is moderate. Here we derive an alternative moment-matching estimator for unrelated subject studies where the common environmental matrix Λ vanishes. Specifically, the multivariate model ***Y = G + E*** and its distributional assumptions vec(*G*) ~ N(0, Σ_*A*_ ⊗ *K*) and vec(*E*) ~ N(0, Σ_*E*_ ⊗ *I*) lead to the following relationship:

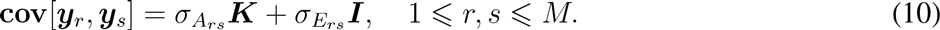

Therefore, an unbiased estimator of *σ_A_rs__* and *σ_E_rs__* can be obtained by regressing 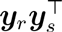, the empirical estimate of the phenotypic covariance matrix cov[*y_r_,y_s_*], onto the the empirical genetic similarity matrix *K* and identity matrix *I*. In particular, we consider the following multiple regression problem:

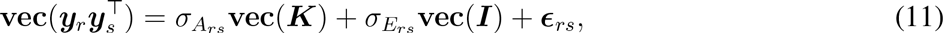

where *∊_rs_* is the residual of this regression. This approach is essentially the Haseman-Elston regression for the classical heritability analysis [59, 60], and has been extended recently to handle various study designs including case-control studies, and more generally termed as phenotype correlation-genetic correlation (PCGC) regression [35]. The ordinary least squares (OLS) estimator of the multiple regression problem (11) satisfies the linear system:

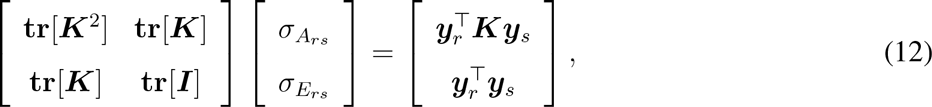

and can be explicitly written

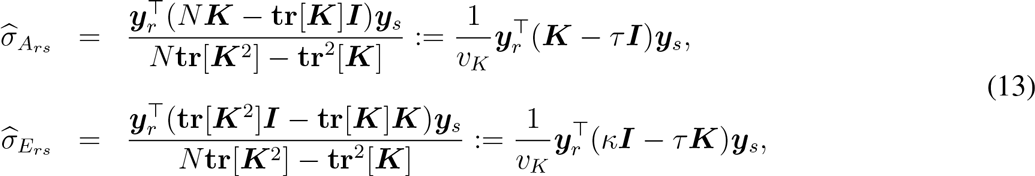

where we have defined τ = tr[*K*]/*N*, κ = tr[*K*^2^]/*N* and *v*_*K*_ = tr[*K*^2^] – tr^2^[*K*]/N = *N*(κ–τ^2^). Therefore, it can be seen that unbiased estimates of the genetic and environmental covariance matrices are as follows:

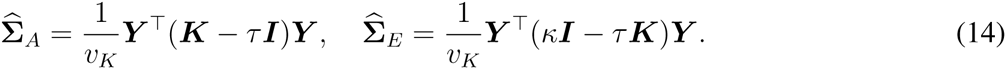

Let 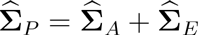, the SNP heritability of a multidimensional trait can then be estimated as

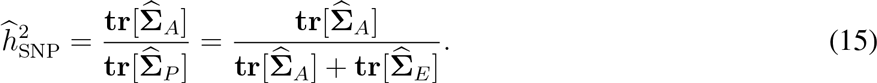

For scalar traits, Eq. (15) degenerates to the classical Haseman-Elston regression estimator.

### Sampling variance of the point estimator

We now derive the variance of 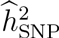. For notational simplicity, we denote *Q*_*A*_ = (*K* – τ*I*)/*v_K_, Q_E_* = (κ*I* – τ*K*)/*v*_*K*_, and also 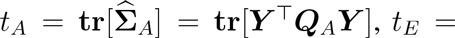 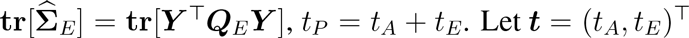. Let *t* = (*t*_*A*_, *t*_*E*_)^T^ and define *f(t) = t_A_/(t_A_ + t_E_) = t_A_/t_P_*. Using a Taylor expansion, we can approximate the variance of the function *f* as follows:

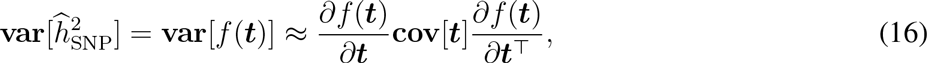

where

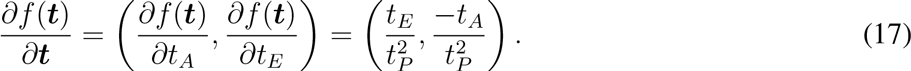

To compute cov [*t*], we define *V*_*rs*_ = cov [*y*_*r*_, *y*_*s*_] = *σ*_*A*_*rs*__ *K* + σ_*E*_*rs*__ *I*, and notice that for any symmetric matrices *Q*_α_ and *Q*_*β*_,

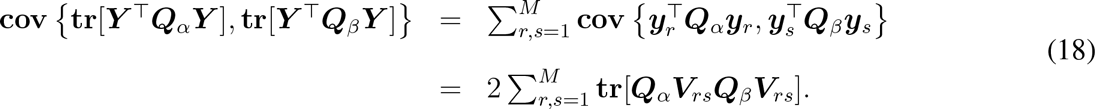

Therefore,

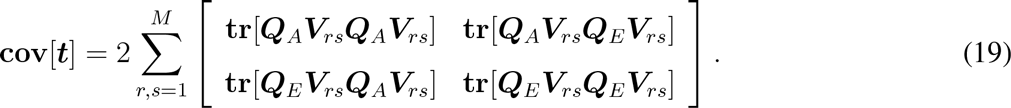

Eq. (19) can be computationally expensive for a moderate or large *M*. We approximate Eq. (19) by using two assumptions that are often made when estimating the sampling variance of the heritability estimator in the study of unrelated individuals [61]: (1) the off-diagonal elements in the empirical genetic similarity matrix *K* are small, such that *K ≈ I* and *V*_*rs*_ = σ_*A*_*rs*__*K* + σ*_E_rs__I* ≈ σ*_A_rs__I* + σ*_E_rs__I* = σ*_P_rs__I*; and (2) the phenotypic covariance Σ_*P*_ is known or can be estimated with very high precision. Using assumption (1), the covariance of *t* can be simplified as follows:

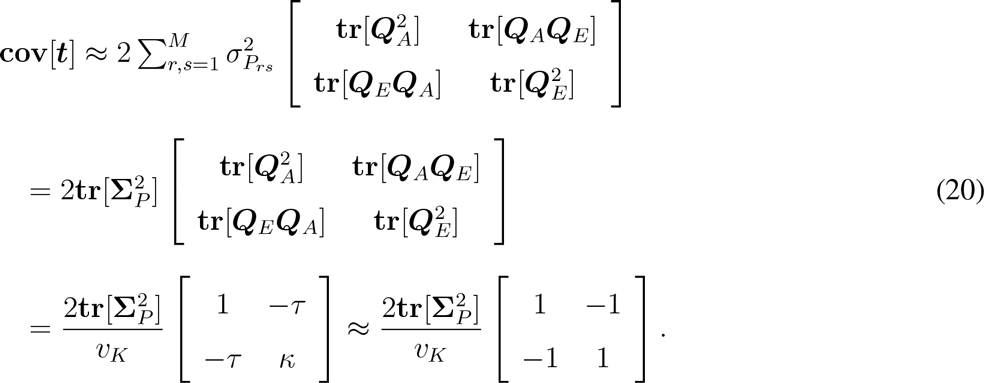

Therefore,

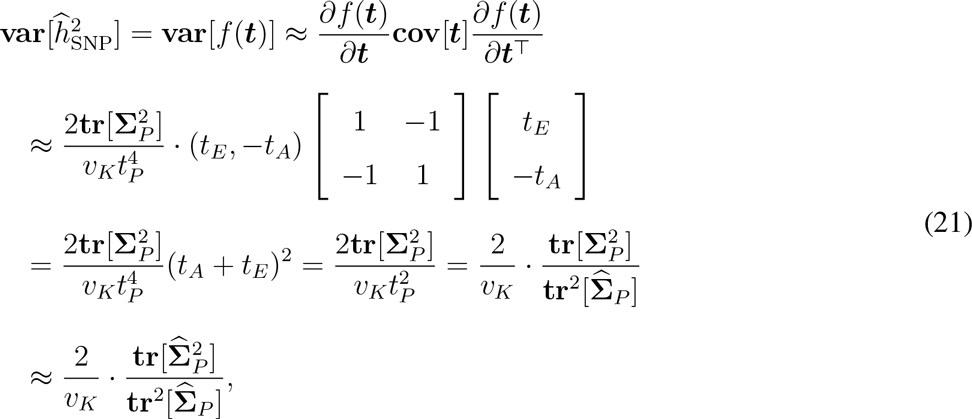

where in the last approximation we have used assumption (2) and replaced Σ_*P*_ with its empirical estimate 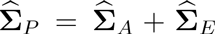. We note that given the empirical genetic similarity matrix *K*, the estimator (21) only depends on the sample size and the phenotypic covariance structure.

For scalar traits, 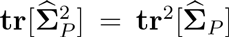, and the estimator (21) degenerates to 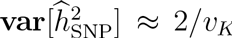, which coincides with existing results in the literature [61]. In general, the covariance matrix 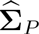 is non-negative definite. Let μ_1_ ≥ μ_2_ ≥ … ≥ μ_M_ ≥ 0 denote its eigenvalues, we have

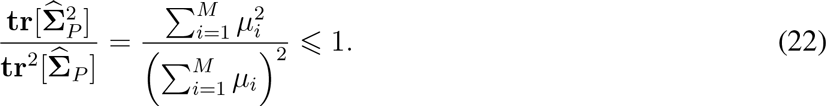

This inequality becomes an equality if and only if 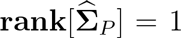, i.e., the *M* traits are perfectly correlated. Therefore, combining multiple traits reduces the variability of heritability estimates relative to analyzing each trait individually.

### Statistic inference

To measure the significance of a heritability estimate, a *p*-value can be computed by conducting a Wald test. Since the null hypothesis, 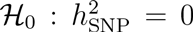, lies on the boundary of the parameter space, the Wald test statistic is distributed as

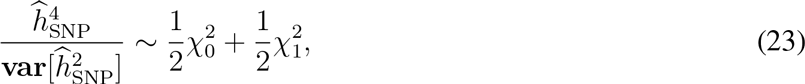

a half-half mixture of 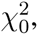, a chi-squared distribution with all probability mass at zero, and 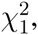, a chi-squared distribution with 1 degrees of freedom [62].

Alternatively, permutation inference can be used by shuffling the rows and columns of the empirical genetic similarity matrix *K*. For each permutation *r* = 1, 2, …, *N*_perm_, we record the heritability estimate 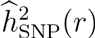 computed from the permuted data. Then for an observed heritability estimate 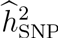, the permutation *p*-value can be computed as [63]

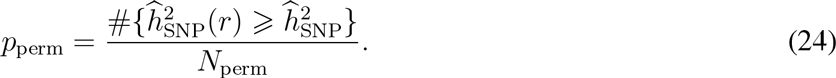

### Heritability estimation in familial studies

A moment-matching estimator can be analogously derived for familial data analysis but has low statistical efficiency due to the strong correlation between the genetic similarity matrix *K* and the common environmental matrix Λ. Therefore, when analyzing the HCP data, we estimate the heritability of each individual component of a multidimensional trait using the restricted maximum likelihood (ReML) algorithm and combine these estimates using a variance-weighted average as derived in Eq. (9).

To estimate the variance of the heritability estimate of a multidimensional trait, we employ a block bootstrapping procedure whereby families are randomly resampled with replacement to produce a bootstrap sample and the heritability is re-estimated. This procedure is repeated for *N*_boots_ times (*N*_boots_ = 1,000 in the present study) to yield bootstrap heritability estimates 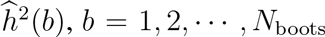. The variance of the heritability estimate is then estimated as [64]

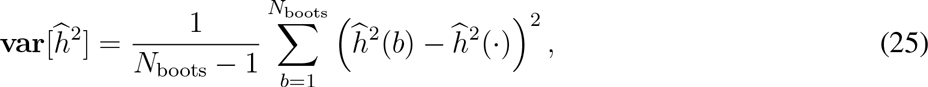

where 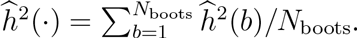

### Modeling covariates

When covariates or nuisance variables need to be adjusted, model (3) becomes a multivariate linear mixed effects model:

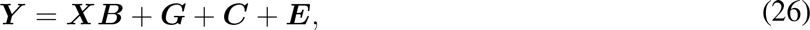

with the distributional assumptions vec(*G*) ~ N(0, Σ_*A*_ ⊗ *K*), vec(*C*) ~ N(0, Σ_*C*_ ⊗ Λ) and vec(*E*) ~ N(0, Σ_E_ Λ *I*), where *X* is an *N × q* matrix of covariates, and *B* is a *q x M* matrix of fixed effects. We employ a strategy described in [65] to removes the covariate matrix from the model. Specifically, there exists an *N x (N – q)* matrix *U*, satisfying *U^T^U = I_(N−q) × (N−q)_*, *UU*^T^ = *P*_0_, and *U*^T^*X* = 0, where *P*_0_ = *I – X*(*X*^T^*X*)^−1^ *X^T^*. The matrix *U*^T^ projects the data from the *N* dimensional space onto an *N – q* dimensional subspace:

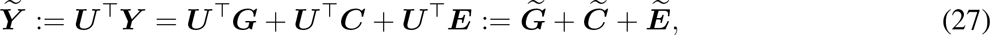

where 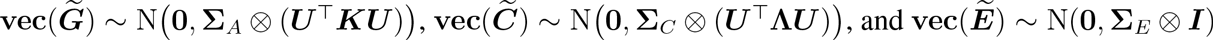. The transformed model is the same as model (3) and thus all estimation and inferential methods developed above can be applied.

### The Brain Genomics Superstruct Project (GSP)

The Harvard/Massachusetts General Hospital (MGH) Brain Genomics Superstruct Project (GSP) is a neuroimaging and genetics study of brain and behavioral phenotypes. More than 3,500 native English-speaking adults with normal or corrected-to-normal vision were recruited from Harvard University, MGH, and the surrounding Boston communities. To avoid spurious effects resulting from population stratification, we restricted our analyses to 1,320 young adults (18-35 years) of non-Hispanic European ancestry with no history of psychiatric illnesses or major health problems (age, 21.54±3.19 years; female, 53.18%; right-handedness, 91.74%). All participants provided written informed consent in accordance with guidelines set by the Partners Health Care Institutional Review Board or the Harvard University Committee on the Use of Human Subjects in Research. For further details about the recruitment process, participants, and imaging data acquisition, we refer the reader to Holmes et al. [36, 37].

### The Human Connectome Project (HCP)

The Human Connectome Project (HCP) collects imaging, behavioral, and demographic data from a large population of healthy adults and aims to shed light on anatomical and functional connectivity within the healthy human brain. We used preprocessed structural MRI data from the WU-Minn HCP consortium and selected subjects that have a similar age range (22-35 years) and ancestry (non-Hispanic/Latino European) as the GSP sample. The 590 subjects we analyzed (age, 29.21±3.45 years; female 55.93%) come from 249 families and comprise 72 monozygotic (MZ) twin pairs, 69 dizygotic (DZ) twin pairs, 253 full siblings of twins and 55 singletons (single birth individuals without siblings). Further details about the recruitment process, imaging data acquisition and MRI data preprocessing can be found in [41, 66].

### Genetic analysis

We used PLINK 1.90 (https://www.cog-genomics.org/plink2) [67], to preprocess the GSP genome-wide SNP data. Major procedures included sex discrepancy check, removing population outliers, spuriously related subjects and subjects with low genotype call rate (< 97%). Individual markers that contained an ambiguous strand assignment and that did not satisfy the following quality control criteria were excluded from the analyses: genotype call rate ≥ 97%, minor allele frequency (MAF) ≥ 1%, and Hardy-Weinberg equilibrium *p* ≥ 1 × 10^−6^. 574,632 SNPs remained for analysis after quality control. We performed a multidimensional scaling (MDS) analysis to ensure that no clear population stratification and outliers exist in the sample (Supplementary Fig. 3). The genetic similarity matrix was estimated from all genotyped autosomal SNPs.

### Laplace-Beltrami Spectrum based shape descriptor

The intrinsic geometry of any 2D or 3D manifold can be characterized by its Laplace-Beltrami Spectrum (LBS) [38, 39], which is obtained by solving the following Laplacian eigenvalue problem (or Helmoltz equation):

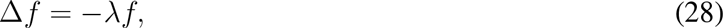

where Δ is the Laplace-Beltrami operator, a generalization of the Laplacian in the Euclidean space to manifolds, *f* is a real-valued eigenfunction defined on a Riemannian manifold, and λ is the corresponding eigenvalue. Eq. (28) can be solved by the finite element method, yielding a diverging sequence of eigenvalues 0 ≤ λ_1_ ≤ λ_2_ ≤ … ↑ + ∞. An implementation of the algorithm is freely available (http://reuter.mit.edu/software/shapedna). The first *M* eigenvalues of the LBS can be used to define a description of the object, which provides a numerical fingerprint or signature of the shape, and is thus known as (length-*M*) “Shape-DNA”.

### Shape analysis pipeline

We used FreeSurfer (http://freesurfer.net) [68], version 4.5.0, a freely available, widely used, and extensively validated brain MRI analysis software package, to process the GSP structural brain MRI scans and label subcortical brain structures. HCP MRI scans were preprocessed by the WU-Minn HCP consortium, and the label files of subcortical structures have been made available [41, 66]. Surface meshes of brain structures were obtained via marching cubes from subcortical segmentations. We created triangular meshes on the boundary surfaces for 20 structures. We then geometrically smoothed these meshes and solved the eigenvalue problem of the 2D Laplace-Beltrami operator on each of these representations, yielding the LBS-based shape descriptor [40]. A python implementation of this pipeline is freely available (http://reuter.mit.edu/software/brainprint).

### Heritability analyses of neuroanatomical shape

We treated the length-*M* LBS-based shape descriptor of each structure as a multidimensional trait and quantified its heritability. In the case of a closed manifold without a boundary, the first eigenvalue is always zero and was thus removed from analysis. Theoretical and empirical evidence have confirmed that the eigenvalues grow linearly and their variance grows quadratically [38, 40]. To avoid that higher eigenvalues dominate the phenotypic covariance, we re-weighted the *m*-th eigenvalue for the *i*-th subject as [38]:

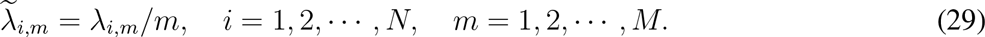

This ensures a balanced contribution of lower and higher eigenvalues on the phenotypic covariance. The LBS also depends on the overall size of the structure. To measure the genetic influences on the shape that are complementary to volume, we further scaled the eigenvalues as:

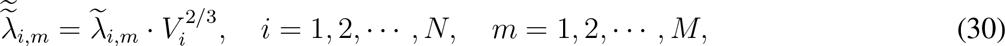

where *V*_*i*_ is the volume of the structure for the *i*-th subject. Since scaling the eigenvalues by a factor *η* results in scaling the underlying manifold by a factor *η*^−1/2^ [38], the normalization (30) ensures that the volumes of the structure are identical across individuals.

We combined the left and right structures in our heritability analyses by averaging their volumetric measurements and concatenating their re-weighted and scaled shape descriptors into one multidimensional trait. We included age, gender, handedness, scanner group, console group, and the top ten principal components of the empirical genetic similarity matrix as covariates when analyzing the GSP sample, and included age, gender and handedness as covariates when analyzing the HCP sample. To remove potential size effect, we always explicitly included the volume of the corresponding structure as a covariate in our shape analyses.

The number of eigenvalues incorporated in the LBS-based shape descriptor and the amount of smoothing applied to the surface mesh are crucial study designs, which might have an impact on heritability estimates. In particular, incorporating a very small number of eigenvalues may be insufficient to characterize the shape of a structure, while very large eigenvalues typically capture fine-scale details, which can be noise and thus might reduce sensitivity. In this study, we reported results obtained by incorporating 50 eigenvalues in the shape descriptor and applying 3 iterations of geometric smoothing to the surface mesh. We conducted sensitivity analyses and confirmed that in the present shape analysis the results were largely robust to different parameter settings (Supplementary Fig. 1).

### Visualizing the principal mode of shape variation

We note that, as shown above, our definition of the heri-tability of a multidimensional trait is a variance-weighted average of individual components, and is invariant to the rotation of the trait vector. Therefore, an equivalent definition of the heritability of a length-*M* LBS-based shape descriptor is the variance-weighted average of the heritability of the first *M* principal components (PCs) of the descriptor, because principal component analysis (PCA) is essentially a rotation of the data. The first PC thus explains the greatest shape variation and has the largest impact on the overall heritability estimate of the shape.

To visualize shape variation along the first PC of the shape descriptor for a given structure, we first aligned the structures from all subjects to a template, *fsaverage*, which is a population average distributed with FreeSurfer [68], using a 7-parameter (global scaling plus 6-parameter rigid body transformation) registration with linear interpolation. Both individual structures and the template were represented with binary label maps, where voxels within the corresponding segmentation label had one and the remainder of the volume had zero values. The registration algorithm maximized the overlap, measured with the Dice score [69], between the corresponding label maps (the fixed template and moving subject which was interpolated and thresholded at 0.5). Note that LBS is invariant to the spatial position and orientation of an object, and we had normalized the shape descriptor for volume in all the analyses. Thus this registration has no impact on the results of our heritability analyses. We then created a sample-specific population average of the structure by computing a weighted average of the interpolated subject images. In particular, each subject was associated with a weight equal to a Gaussian kernel centered around the mean of the first PC and evaluated at the subject’s first PC of the shape descriptor. The width of the kernel was selected such that 500 shapes received non-zero weights. The isosurface of the resulting probability map at 0.5 was used to represent the average shape of the structure, and all visualizations were presented on this surface.

The same Gaussian kernel was used to generate average probability images for shapes centered at the two extremes (±2 standard deviation or SD) of the principal axis. These average probability images were offset to achieve identical volumes when thresholded at 0.5. The difference of the two extreme shapes were depicted on the sample-specific population average, by visualizing the difference in the probability values. Blue indicated that the average shape at −2 SD achieved a higher probability value and thus was larger in those regions than the average shape at the +2 SD. For red regions, the opposite was true.

### Data availability

The Brain Genomics Superstruct Project (GSP) data analyzed during the current study are publicly available at http://neuroinformatics.harvard.edu/gsp/. The Human Connectome Project (HCP) data analyzed during the current study are publicly available at http://www.humanconnectome.org. Other data are available from the corresponding author on reasonable request.

## Acknowledgements

This research was carried out in whole at the Athinoula A. Martinos Center for Biomedical Imaging at the Massachusetts General Hospital (MGH), using resources provided by the Center for Functional Neuroimaging Technologies, P41EB015896, a P41 Biotechnology Resource Grant supported by the National Institute of Biomedical Imaging and Bioengineering (NIBIB), National Institutes of Health (NIH). This work also involved the use of instrumentation supported by the NIH Shared Instrumentation Grant Program; specifically, grant numbers S10RR023043 and S10RR023401.

Data were provided in part by the Brain Genomics Superstruct Project (GSP) of Harvard University and MGH, with support from the Center for Brain Science Neuroinformatics Research Group, Athinoula A. Martinos Center for Biomedical Imaging, Center for Human Genetic Research, and Stanley Center for Psychiatric Research. 20 individual investigators at Harvard and MGH generously contributed data to the overall project.

Data were also provided in part by the Human Connectome Project, WU-Minn Consortium (Principal Investigators: David Van Essen and Kamil Ugurbil; 1U54MH091657) funded by the 16 NIH Institutes and Centers that support the NIH Blueprint for Neuroscience Research; and by the McDonnell Center for Systems Neuroscience at Washington University.

This research was also funded in part by NIH grants K25CA181632 (to MR); K01MH099232 (to AJH); K99MH101367 (to PHL); R01NS083534, R01NS070963, and 1K25EB013649-01 (to MRS); K24MH094614 and R01MH101486 (to JWS); an MGH ECOR Tosteson Postdoctoral Fellowship Award (to TG); the Brazilian National Research Council (CNPq), grant number 211534/2013-7 (to AMW); and a BrightFocus Foundation grant AHAF-A2012333 (to MRS). JWS is a Tepper Family MGH Research Scholar.

## Competing interests

The authors declare no competing financial interests.

